# CIGAR-seq, a CRISPR/Cas-based method for unbiased screening of novel mRNA modification regulators

**DOI:** 10.1101/2020.10.03.324715

**Authors:** Liang Fang, Wen Wang, Guipeng Li, Li Zhang, Jun Li, Diwen Gan, Jiao Yang, Yisen Tang, Zewen Ding, Min Zhang, Wenhao Zhang, Daqi Deng, Zhengyu Song, Qionghua Zhu, Huanhuan Cui, Yuhui Hu, Wei Chen

**Author notes:** These authors contributed equally to this work (Liang Fang, Wen Wang, Guipeng Li).

## Abstract

Cellular RNA is decorated with over 170 types of chemical modifications. Many modifications in mRNA, including m^6^A and m^5^C, have been associated with critical cellular functions under physiological and/or pathological conditions. To understand the biological functions of these modifications, it is vital to identify the regulators that modulate the modification rate. However, a high-throughput method for unbiased screening of these regulators is so far lacking. Here, we report such a method combining pooled CRISPR screen and reporters with RNA modification readout, termed <u>C</u>RISPR <u>i</u>ntegrated <u>g</u>RNA <u>a</u>nd <u>r</u>eporter sequencing (CIGAR-seq). Using CIGAR-seq, we discovered NSUN6 as a novel mRNA m^5^C methyltransferase. Subsequent mRNA bisulfite sequencing in HAP1 cells without or with NSUN6 and/or NSUN2 knockout showed that NSUN6 and NSUN2 worked on non-overlapping subsets of mRNA m^5^C sites, and together contributed to almost all the m^5^C modification in mRNA. Finally, using m^1^A as an example, we demonstrated that CIGAR-seq can be easily adapted for identifying regulators of other mRNA modification.

## Introduction

Cellular RNAs can be chemically modified in over a hundred different ways, and such modifications have been associated with diverse cellular functions under physiological and/or pathological conditions (Machnicka, Milanowska et al., 2013, Roundtree, Evans et al., 2017). To achieve so called dynamic epitranscriptomic regulation, each modification needs its distinct deposition, removal and recognition factors (termed ‘writers’, ‘erasers’ and ‘readers’, respectively). Yet, comparing to the ever-expanding techniques on detecting RNA modifications (Zhao, Song et al., 2020), the methods to systematically identify writers and erasers of RNA modifications are rather limited. For instance, the first N6-adenosine (m^6^A) methyltransferase METTL3 was identified through a combination of in vitro assay, conventional chromatography, electrophoresis and microsequencing (Bokar, Rath-Shambaugh et al., 1994, Bokar, Shambaugh et al., 1997), and the second m^6^A methyltransferase METTL14 was discovered through the phylogenetic analysis based on METTL3 (Liu, Yue et al., 2014). In general, the first strategy is less efficient and may have assay-specific bias, while the second strategy relies on the prior knowledge of related molecule(s). So far, unbiased method to screen for novel regulators of RNA modifications is still lacking.

Recently, the rapid development of CRISPR-based gene manipulation provides a new paradigm for high-throughput and genome-wide functional screening. Pooled CRISPR screen outperforms array-based screen by its scalability and low cost, however, was largely restricted to standard readouts, including survival, proliferation and FACS-sortable markers (Hanna & Doench, 2020). Most recently, combining with microscopy-based approaches, CRISPR screen enabled the association of subcellular phenotypes with perturbation of specific gene(s) (Wheeler, Vu et al., 2020, Yan, Stuurman et al., 2020). in studying regulation of gene expression, Perturb-seq, CRISP-seq and CROP-seq, which combine CRISPR-based gene editing with single-cell mRNA sequencing, allowed transcriptome profile to serve as comprehensive molecular readout (Adamson, Norman et al., 2016, Datlinger, Rendeiro et al., 2017, Dixit, Parnas et al., 2016), but often with limited throughput. Until now, pooled CRISPR screen with epitranscriptomic readout has not yet been developed.

One important RNA modification, 5-methylcytosine (m^5^C), was first identified in stable and highly abundant tRNA and rRNA (Agris, 2008, Helm, 2006, Schaefer, Pollex et al., 2009). Subsequently, many novel m^5^C sites in mRNA were discovered by using next-generation sequencing-based methods, including mRNA bisulfite sequencing (mRNA-BisSeq) (Schaefer et al., 2009, Squires, Patel et al., 2012), m^5^C-RNA immunoprecipitation (RIP) (Edelheit, Schwartz et al., 2013), 5-azacytidine-mediated RNA immunoprecipitation (Aza-IP) (Khoddami & Cairns, 2013) and methylation-individual-nucleotide-resolution crosslinking and immunoprecipitation (miCLIP) (Hussain, Sajini et al., 2013). The m^5^C modification has been reported to regulate the structure, stability and translation of mRNAs (Guallar, Bi et al., 2018, Li, Li et al., 2017, Luo, Feng et al., 2016, Schumann, Zhang et al., 2020, Shen, Zhang et al., 2018), and be catalyzed by NOP2/Sun RNA methyltransferase family member 2 (NSUN2) (David, Burgess et al., 2017, Khoddami & Cairns, 2013, Yang, Yang et al., 2017a). However, recent studies have shown that, even after NSUN2 knockout (KO), a significant number of m^5^C sites in mRNA remained methylated (Huang, Chen et al., 2019, Trixl & Lusser, 2019), suggesting the existence of additional methyltransferase(s) involved in mRNA m^5^C modification. To fully appreciate the function of this modification, it would be important to identify the remaining methyltransferase(s).

Here, we report a method combining pooled CRISPR screen and a reporter with epitrancriptomic readout, termed <u>C</u>RISPR <u>i</u>ntegrated <u>g</u>RNA <u>a</u>nd <u>r</u>eporter sequencing (CIGAR-seq). Using CIGAR-seq with a reporter containing a m^5^C modification site, we screening through a gRNA library targeting 829 RNA-binding proteins and identified NSUN6 as a novel m^5^C writer of mRNA. mRNA-BisSeq in HAP1 cells without or with NSUN6 and/or NSUN2 knockout showed NSUN6 and NSUN2 worked on non-overlapping subsets of mRNA m^5^C sites, and together contributed to almost all the m^5^C modification in mRNA. Finally, using m^1^A as an example, we demonstrated that CIGAR-seq can be easily adapted for studying other mRNA modification.

## Results

### CIGAR-seq: Pooled CRISPR screening with a epitranscriptomic readout

In CIGAR-seq, to integrate pooled CRISPR screening with a epitranscriptomic readout, here more specifically m^5^C modification readout, we adopted the previously developed CROP-seq method (Datlinger et al., 2017) and replaced the WPRE cassette on the original vector by an endogenous m^5^C site with its flanking region (Fig 1A). Thereby, the mRNA molecules transcribed from this lentiviral vector contain a selection marker followed by an endogenous m^5^C site, a U6 promoter and a gRNA sequence. To detect the m^5^C level in the gRNA sequence-containing transcripts, total mRNA was firstly subjected to bisulfite treatment followed by reverse transcription. Subsequently, a primer pair flanking the m^5^C site and gRNA sequence was used to amplify the region for Sanger or next-generation sequencing (Fig 1A). in this way, the methylation level of the m^5^C reporter site can be measured and associated with the gRNA targeting a specific gene.

**Figure 1.**
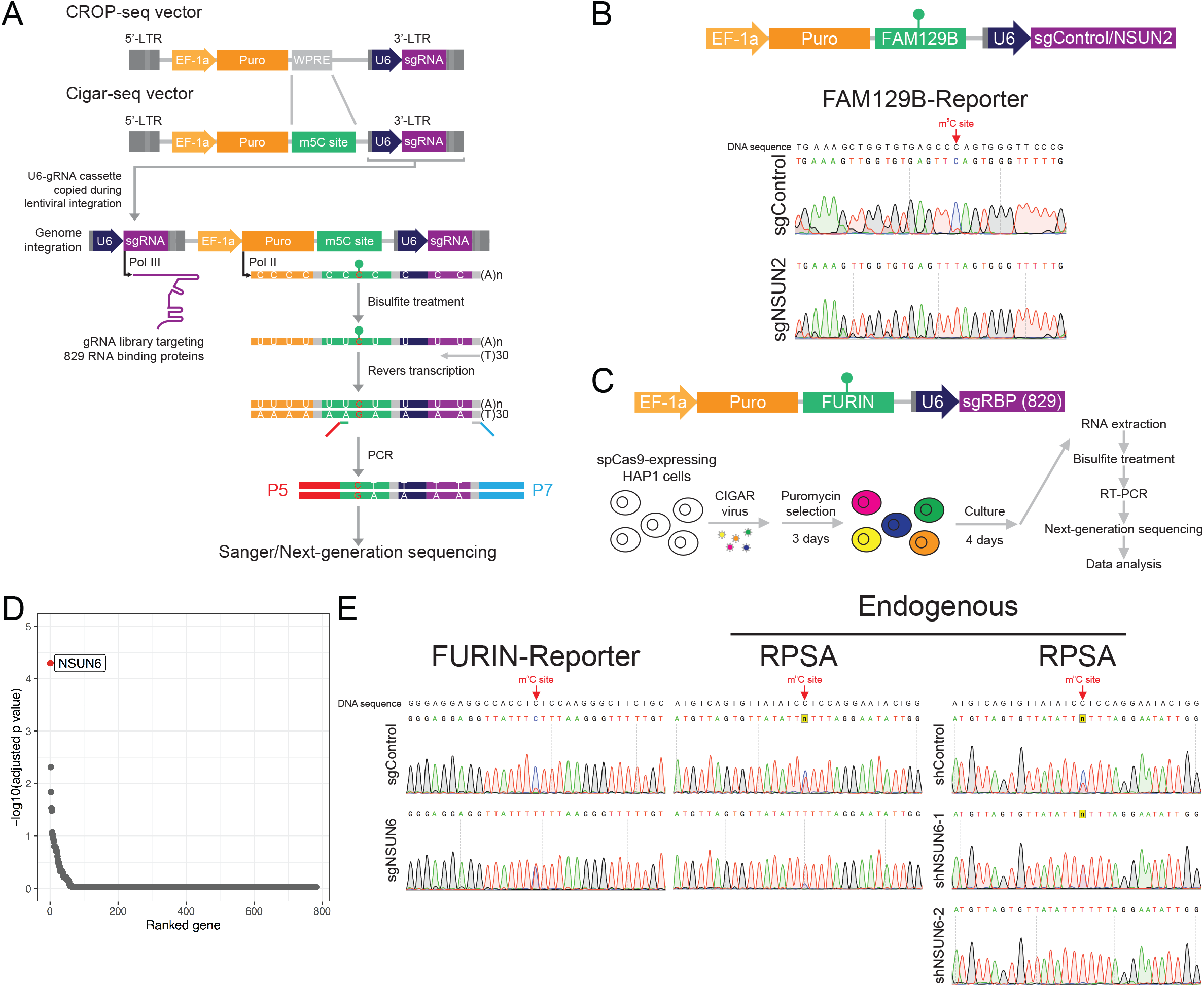
CIGAR-seq identified NSUN6 as a novel mRNA methyltransferase. **A**, An illustration of CIGAR-seq method in studying m^5^C modification. The WPRE cassette on the original CROP-seq vector was replaced by an endogenous m^5^C site with its flanking region. To measure the m^5^C level of the reporter site in gRNA sequence-containing transcripts, mRNA was subjected to bisulfite treatment followed by reverse transcription. Then, a primer pair flanking the m^5^C site and gRNA sequence was used to amplify the region for subsequent Sanger or next-generation sequencing. **B**, Validation of CIGAR-seq method using a m^5^C reporter site derived from FAM129B gene with a control gRNA without target genes and a gRNA targeting NSUN2. Upon NSUN2 knockout, the m^5^C modification is diminished in the FAM129B reporter mRNA, whereas the modification remains intact in control knockout cells. **C**, The application of CIGAR-seq in screening for regulators of m^5^C sites. Cas9-expressing HAP1 cells were transduced with viral particles that express Cigar vectors combining NSUN2-independent m^5^C reporter sites derived from FURIN gene and a gRNA library targeting 829 RBPs. Seven days after transduction, enriched polyA RNA was bisulfite-treated, reveres-transcribed and PCR-amplified using primers flanking the m^5^C site and gRNA sequence to generate NGS library. **D**, The rank of gene whose knockout reduced m^5^C modification rate of the reporter site (Methods). NSUN6 was the top hit. **E**, Validation of NSUN6 as a mRNA m^5^C methyltransferase. Knockout as well as knockdown NSUN6 reduced the m^5^C level in both FURIN m^5^C reporter transcripts and endogenous NSUN2-independent m^5^C sites in RPSA gene.

As a proof-of-concept experiment, a known NSUN2-dependent m^5^C site in FAM129B (also known as NIBAN2) gene was cloned into CIGAR-seq vector, together with a control gRNA without any target gene or a gRNA targeting NSUN2 (Fig 1B, upper panel). Seven days after transduction into Cas9-expressing HAP1 cells, m^5^C modification level on the reporter transcripts was measured. The result demonstrated that, upon NSUN2 perturbation (Fig EV1A), the m^5^C modification rate reduced significantly in the reporter containing NSUN2 targeting gRNAs (Fig 1B, lower panel), whereas the modification remained intact in the reporter containing the control gRNA (Fig 1B, upper panel).

### CIGAR-seq identified NSUN6 as a novel mRNA methyltransferase

As suggested by previous studies that large amount of m^5^C sites in mRNA remained methylated after NSUN2 knockout (Huang et al., 2019, Trixl & Lusser, 2019), we sought to utilize CIGAR-seq to identify gene(s) that mediate the m^5^C modification on NSUN2-independent sites. First, to determine the NSUN2-independent m^5^C sites, we established NSUN2 knockout (NSUN2-KO) HAP1 cells (Fig EV4A), and performed mRNA bisulfite sequencing (mRNA-BisSeq) in wildtype as well as NSUN2-KO cells. Following bioinformatic pipeline proposed by Huang et al. (Huang et al., 2019), a set of 208 m^5^C sites was identified in wildtype HAP1 cells (Methods), only 90 (43.3%) of which showed significantly reduced m^5^C level in NSUN2-KO cells (Fig EV1B).

We then chose a NSUN2-independent site in the 3’UTR of FURIN gene with a high m^5^C modification rate as the reporter site for CIGAR-seq. Meanwhile, a gRNA library targeting 829 RNA binding proteins (RBP) was synthesized (Table EV1). To establish the CIGAR-seq vector pool for the genetic screen, the gRNA library was firstly cloned into the vector followed by the insertion of the FURIN m^5^C site with its flanking genomic region (Methods). Cas9-expressing HAP1 cells were then transduced with CIGAR-seq virus pool (Fig 1C, Methods). Seven days after transduction, cells were collected and subjected to RNA extraction. Enriched polyA RNA was then bisulfite-treated, reveres-transcribed and PCR-amplified using primers flanking the m^5^C site and gRNA sequence to generate next-generation sequencing (NGS) library (Methods). After pair-end sequencing and data processing, 811 genes were detected with at least one gRNA, of which 782 genes had at least two gRNAs (Fig EV1C). The m^5^C modification rate of reporter site was calculated for each gRNA. While the median m^5^C modification rates of gRNA-associated reporter sites were around 93.5%, a small part of gRNAs showed significantly reduced m^5^C rates (Fig EV1D). To prioritize the candidate genes, Stouffer’s method was used to calculate the combined P value based on the gRNAs targeting the same gene (Methods). As shown in Fig 1D, it turned out that NSUN6, a member of NOL1/NOP2/sun domain (NSUN) family, was identified as the best hit (Fig 1D). Interestingly, NSUN6 was previously reported to introduce the m^5^C in tRNA (Li, Li et al., 2019). However, two previous studies did not show significant m^5^C changes in mRNA after NSUN6 perturbation in HeLa cells (Huang et al., 2019, Yang et al., 2017a).

To validate our result, a gRNA targeting NSUN6 was inserted into the CIGAR-seq vector containing the FURIN m^5^C site. As shown in Figure 1E, perturbation of NSUN6 (Fig EV2A) indeed reduced the m^5^C level in both reporter mRNA and endogenous NSUN2-independent m^5^C site in RPSA gene (Fig 1E, left and middle panels). Furthermore, to rule out the potential off-target effect of gRNA, we repressed NSUN6 expression using shRNA (Fig EV2B). Again, the m^5^C level at the endogenous site was also reduced in cells with NSUN6 repression (Fig 1E, right panel). Together, these results confirmed NSUN6 as a bona fide mRNA m^5^C methyltransferase.

### Global profiling of NSUN6-dependent m^5^C sites

To globally characterize NSUN6-dependent m^5^C sites, we established NSUN6 knockout (NSUN6-KO) HAP1 cells (Fig EV4B) and performed mRNA-BisSeq. Of 208 m^5^C sites identified in wildtype HAP1 cells, 65 (31.2%) showed significant reduction at m^5^C level in NSUN6-KO cells (Fig 2A). To illustrate the features of sequence flanking NSUN6-dependent m^5^C sites in HAP1 cells, motif analysis was performed based on the upstream and downstream 10 nucleotide sequences flanking the m^5^C sites. As shown in Figure 2B, NSUN6-dependent m^5^C sites were embedded in slightly GC-rich environments with a strongly enriched TCCA motif at 3’ of m^5^C sites. Previously, a similar 3’ TCCA motif was also found at NSUN6 target sites in tRNAs (Li et al., 2019), and has also been proposed as sequence motif around NSUN2-independent sites in another study (Huang et al., 2019). In comparison, the sequence feature around NSUN2-dependent m^5^C sites is distinct, which is enrich for 3’ NGGG motif (Huang et al., 2019, Yang, Yang et al., 2017b).

**Figure 2.**
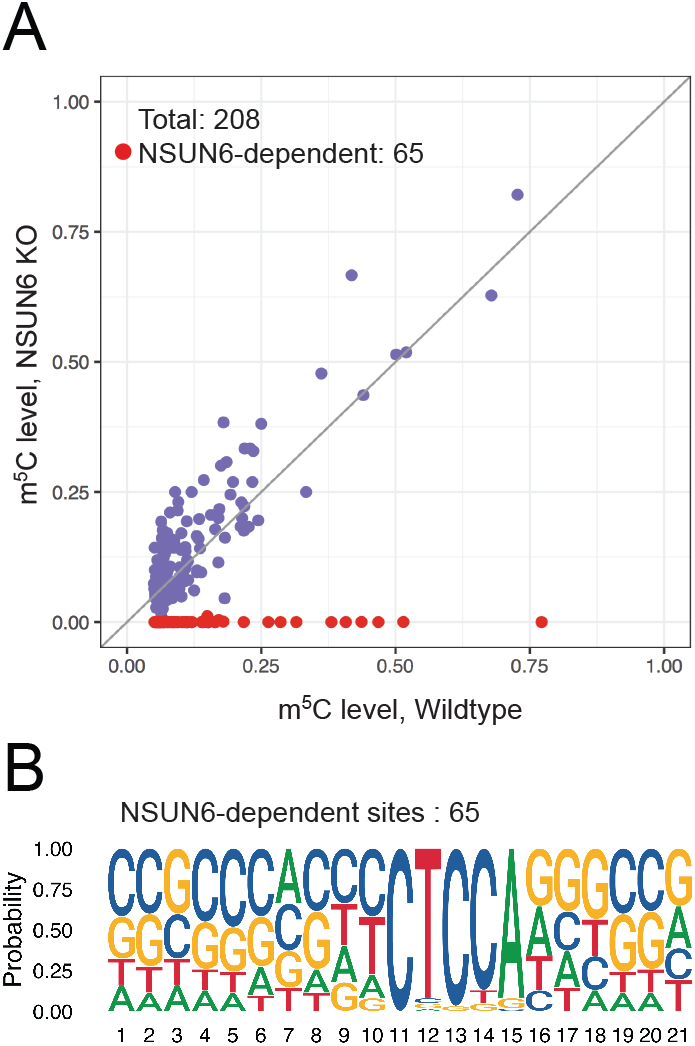
Global profiling of NSUN6-dependent m^5^C sites. **A**, mRNA bisulfite sequencing revealed NSUN6-dependent m^5^C sites in HAP1 cells. Of 208 m^5^C sites identified in wildtype cells, 65 showed significantly reduced modification in NSUN6 knocked out cells. X and Y axis represented the modification rate in wildtype and NSUN6 knocked out HAP1 cells, respectively. **B**, The sequence features of NSUN6-dependent m^5^C sites in HAP1 cells. A strong 3’ TCCA motif was found in NSUN6-dependent sites.

### Contribution of NSUN6 and NSUN2 to the mRNA m^5^C modification

We then evaluated the relative contribution of NSUN6 and NSUN2 to the global mRNA m^5^C modification. First, comparing between NSUN2- and NSUN6-dependent m^5^C sites, as shown in Figure 3A, these sites were largely non-overlapping, suggesting their non-redundant biological functions. Then, to further examine whether NSUN2 and NSUN6 together are responsible for all mRNA m^5^C modifications, NSUN2 and NSUN6 double KO (NSUN2/6-dKO) HAP1 cells were established (Fig EV4C) and subjected to mRNA-BisSeq analysis. As shown in Figure 3B, the modification of m^5^C sites depend only on NSUN6 or NSUN2 (62 and 87, respectively) were also abolished in NSUN2/6-dKO cells. While NSUN6-dependent sites were strongly enriched for 3’ TCCA motif as shown earlier, NSUN2-dependent sites were enriched for 3’ NGGG motif as previously reported (Huang et al., 2019, Yang et al., 2017b) (Fig 3C). Furthermore, we carefully examined the three sites that showed dependence on both NSUN6 and NSUN2, as well as the 56 sites that were independent of both NSUN6 and NSUN2. Comparing to the other three groups, the group of three overlapping sites had very low m^5^C level (Fig 3D). In addition, the m^5^C sites in ANGEL1 and ZNF707 possessed a 3’ TCCA and a 3’ AGGG motif, respectively (Fig EV3A&B), suggesting they are very likely a NSUN6- and a NSUN2-dependent site, respectively, but with low m^5^C level that led to false negative findings in the mRNA-BisSeq analysis of some but not all the samples. The remaining m^5^C site in STRN4 was embedded within a cluster of “pseudo” m^5^C sites (Fig EV3C), which was highly likely an artifact due to the incomplete bisulfite conversion as suggested before (Haag, Warda et al., 2015, Huang et al., 2019). Similarly, the group of 56 NSUN2/6-independent sites were also highly enriched for such clusters of pseudo m^5^C sites: 52 sites had at least one pseudo m^5^C site in vicinity (Table EV2). The remaining four sites all had very low m^5^C level.

**Figure 3.**
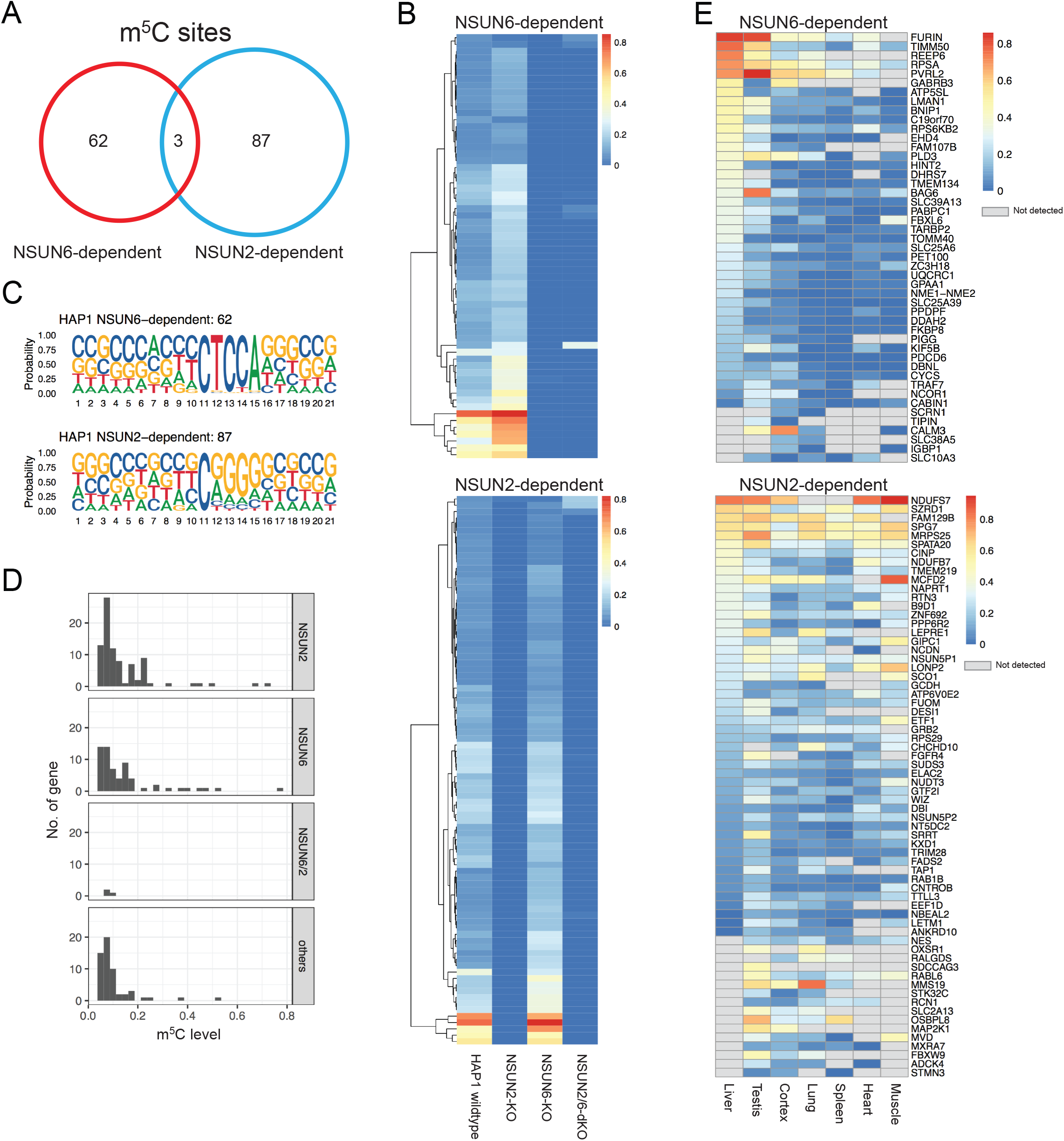
Comparison between NSUN6- and NSUN2-dependent m^5^C modification sites. **A**, The largely non-overlap between NSUN2- and NSUN6-dependent m^5^C sites. **B**, Heatmap showing the m^5^C modification rate in wildtype, NSUN2-KO, NSUN6-KO and NSUN2/6-dKO HAP1 cells for NSUN6- (upper panel) and NSUN2-dependent sites (lower panel), respectively. **C**, The sequence features of NSUN6-only- (upper panel) and NSUN2-only-dependent m^5^C sites (lower panel) in HAP1 cells. While NSUN6-dependent sites were strongly enriched for 3’ TCCA motif, NSUN2-dependent sites were enriched for 3’ NGGG motif. **D**, The modification rate of 4 groups of m^5^C sites that showed different dependence. Comparing to the other three groups, the group of three overlapping sites showed very low m5C level. **E**, Modification rate of NSUN6- and NSUN2-dependent m5C sites across different tissues.

To explore the modification rate of NSUN6/2-dependent m^5^C sites across different tissues, we resorted to mRNA-BisSeq data from a previous study (Huang et al., 2019). As shown in Figure 3E, m^5^C modification on 47 NSUN6- and 66 NSUN2-dependent m^5^C sites could also be observed in other human tissue(s). While the modification rate of NSUN6-dependent sites was by and large highest in liver, NSUN2-dependent ones did not show such tissue biases (Fig 3E).

### CIGAR-seq could be used for the study of other mRNA modification

Finally, to explore the potential application of CIGAR-seq in the study of other mRNA modifications, we turned to N-1-methyladenosine (m^1^A). As *N*-1-methyladenosine (m^1^A) can cause mis-incorporation during cDNA synthesis (Hauenschild, Tserovski et al., 2015), its modification can be detected by direct cDNA sequencing. Similar as the previous NSUN2 proof-of-concept experiment, we chose a well-characterized m^1^A site from MALAT1, which is known to be modified by TRMT6/TRMT61A complex (Dominissini, Nachtergaele et al., 2016, Li, Xiong et al., 2017, Safra, Sas-Chen et al., 2017). We cloned the site and its flanking region into CIGAR-seq vector with a control gRNA and two gRNAs targeting TRMT6/TRMT61A complex, respectively (Fig 4A). As shown in Figure 4C, in HAP1 cells with perturbation of either TRMT6 or TRMT61A (Fig 4B), the m^1^A modification of the reporter site was completely abolished, whereas in cells transduced with control gRNAs, the modification remains intact.

**Figure 4.**
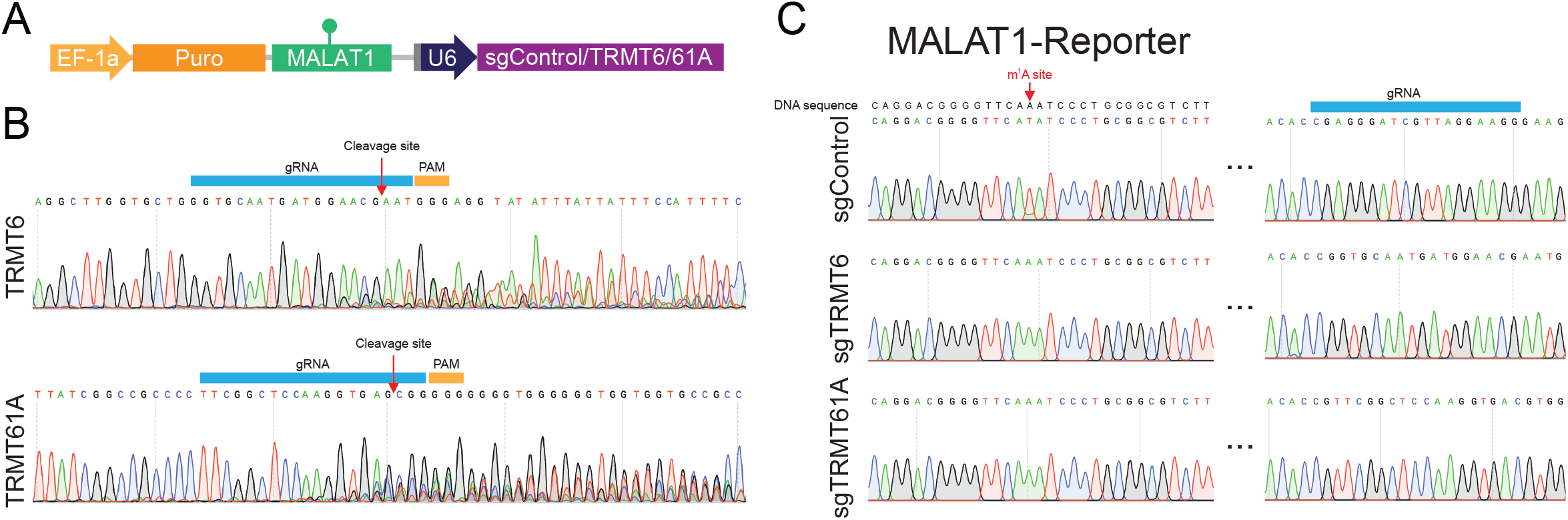
Exemplar application of CIGAR-seq in the study of m^1^A modification. **A,** An illustration of CIGAR-seq vector designed for m^1^A modification. A known TRMT6/61A complex-dependent m^1^C site in MALAT1 gene was cloned into CIGAR-seq vector, together with control gRNA as well as gRNAs targeting TRMT6 and TRMT61A, respectively. **B&C**, Upon perturbation of either TRMT6 or TRMT61A in HAP1 cells, the m^1^A modification was completed abolished in m^1^A reporter site, whereas the modification remains intact in control knockout cells.

## Discussion

Combining pooled CRISPR screening strategy and a reporter with epitranscriptomic readout, CIGAR-seq for the first time enables the unbiased screening for novel regulators of mRNA modifications. In this study, we demonstrated its power in identification of NSUN6 as a novel mRNA m^5^C methyltransferase. in addition, we also showed its potential application in studying m^1^A modification. Integrating additional modification readout strategies into our pipeline, it could be further adapted to investigate other modifications. For instance, we can use CIGAR-seq to search for potential regulators of RNA editing by simply reading the A-G or C-T changes in the cDNA sequence reads derived from A-I or C-U RNA editing reporters. m^6^A-iCLIP (Linder, Grozhik et al., 2015) or SELECT (Xiao, Wang et al., 2018) method, which were used to measure the modification rate of individual m^6^A site, could also be integrated into our CIGAR-seq in analyzing m^6^A regulators. Furthermore, changing reporters to those with other regulatory readout, for example alternative splicing or alternative polyadenylation pattern, the potential application of CIGAR-seq could be easily extended to screen for factors involved in diverse post-transcriptional regulations. There, given the readout is based on directly measuring the reporter-derived RNAs, CIGAR-seq would be in principle superior to current fluorescence-reporter-FACS based screening strategies.

Like any other high-throughput assays, CIGAR-seq has also its own sensitivity and specificity issues, which could be affected by the choice of reporter and CRISPR system. The reporter could affect its performance in two ways. First, the high or low modification level of the reporter site could result in biased performance in detecting positive or negative regulators. For example, in this study, our m^5^C site from FURIN genes has a very high modification rate. At this level, it would be much more sensitive in finding the decrease of methylation rate, therefore be much easier to discover the methyltransferase than potential demethylase if any in this case. In contrast, the use of reporter site with low modification rate would not be preferable in identifying methyltransferase. Second, except writer and eraser, most regulators may modulate the modification level through binding to the cis-regulatory elements, which are not necessarily in direct vicinity of the target site. A reporter construct with a limited length might not be able to include all the relevant cis-elements. Consequently, we would fail to identify the regulators with binding sites missed in the reporter. On the other hand, the CIGAR-seq vector itself may contain artificial regulatory sequences affecting the modification of reporter site, which could result in the assay-specific artifacts. Therefore, subsequent careful validation with endogenous sites would be essential when working with the CIGAR-seq. The choice of CRISPR system could also have an effect. The screening based on CRISPR/Cas9 system, as applied in this study, would have limitations in finding potential regulators that are essential for cell survival and/or proliferation (e.g. METTL3/METTL14) since the gRNAs targeted at those essential genes would be largely depleted in the final sequencing library. This problem could be potentially alleviated by adopting CRISPRi or CRISPRa systems. In the future, with further improvements in CRISPR system and development of more sequencing-based readout with high precision, CIGAR-seq will become a versatile tool for systematic discoveries of players in multiple layer of RNA-based post-transcriptional gene regulation.

## Methods

### Experimental methods

#### Cell culture and gene manipulation

HAP1 cell was obtained from Horizon discovery and cultured in RMPI1640 medium (22400089, Gibco) with 10% FBS (10270106, Gibco) and 1% P/S (15070063, Gibco) at 37°C with 5% CO2. Cas9-expressing HAP1 cell line was established by using lentiCas9-Blast plasmid (#52962, Addgene). To generate NSUN2-KO, NSUN6-KO and NSUN2/6-dKO clonal HAP1 cells, Cas9-expressing HAP1 cells were transduced with CROP-seq (Addgene, #86708) virus expressing following gRNAs: gNSUN2, 5’-GCTGTTCGAGCACTACTACC-3’; gNSUN6, 5’-GACCTTCAAGATGTGTTACT-3’. NSUN6 knockdown mediated by shRNA was performed using pLKO.1-blast plasmid (modified from pLKO.1-puro, #10878, Addgene) with following shRNAs: shControl, 5’-CGTCTGGCTAATAAGGACTCT-3’; shNSUN6-1, 5’-GCAAAGAAATCTTCAGTGGAT-3’; shNSUN6-2, 5’-GCTGGAGATGTTATTTCTGTA-3’.

#### RT-qPCR

Total RNA was extracted by TRIzol^®^ Reagent (Ambion). First-stand cDNA was synthesized using HiScript III 1^st^ Stand cDNA Synthesis Kit (Vazyme, #R312-02). Quantitative PCR was performed by Hieff qPCR SYBR Green Master Mix (Yeasen, #11201ES08) and the BIO-RAD real-time PCR system. Following primers were used to detect relative gene expression: NSUN6-F, 5’-GGAGCCAAAGAATTTGATGGAACA-3’; NSUN6-R, 5’-ATGCCCATGCCTTTCAGTTC-3’; GAPDH-F, 5’-AGCCACATCGCTCAGACAC-3’; GAPDH-R, 5’-GCCCAATACGACCAAATCC-3’.

#### CIGAR-seq vector with m^5^C/m^1^A reporters and individual gRNA

Sequence flanking m^5^C site of FAM129B was amplified by forward primer 5’-GCCTGAACGCGTTAAGTCGAC-GGCTGGACACTGCTGGGG-3’ and reverse primer 5’-GTAAGTCATTGGTCTTAAAGTCGAC-GGGGAAAGCGAGGCTCG-3’ from genomic DNA; m^5^C site of FURIN by forward primer 5’-GCCTGAACGCGTTAAGTCGAC-CCGGCCCCAGCCAGAGTTC-3’ and reverse primer 5’-GTAAGTCATTGGTCTTAAAGTCGAC-TGGTGGAGGCACGGAGCACA-3’, and m^1^A site of MALAT1 by forward primer 5’-GCCTGAACGCGTTAAGTCGAC-CTTCAGTAGGGTCATGAAGGTTTTTCT-3’ and reverse primer 5’-GTAAGTCATTGGTCTTAAAGTCGAC-ATACATCAAGGATGTATATAGTTCAAAGATATTGTGC-3’.

Amplified products were used to replace WPRE cassette in CROP-seq vector (Addgene, #86708) by ClonExpress II One Step Cloning Kit (Vazyme). Afterwards, following gRNAs were inserted at BsmBI sites to knockout individual genes: gControl, 5’-GAGGGATCGTTAGGAAGGG-3’; gNSUN2, 5’-GCTGTTCGAGCACTACTACC-3’; gNSUN6, 5’-GACCTTCAAGATGTGTTACT-3’; gTRMT6, 5’-GGTGCAATGATGGAACGAAT-3’; gTRMT61A, 5’-TTCGGCTCCAAGGTGACGTG-3’.

#### m^5^C detection by bisulfite conversion followed by sanger sequence

Total RNA was extracted by TRIzol^®^ Reagent (Ambion). mRNA was enriched using VAHTS mRNA Capture Beads (Vazyme, #N401). 200 ng mRNA was converted by EZ RNA methylation kit (Zymo Research) according to the manufacturer’s protocol with minor modification. More specifically, mRNA was incubated at 70 °C for 10 min, and 60 °C for 1 h. Converted RNA was then reverse transcribed into cDNA using HiScript II Q Select RT SuperMix (Vazyme, #R233-01).

To measure m^5^C rate in FURIN m^5^C reporter, target site was amplified using vector specific primer pair 5’-TTGTAATTTTTTTTTTTATGAGTGGTTTGGTTTTA-3’ and 5’-TTAAAAAATAACTAAAATCTACAACTACCTTATAAATCATTAATCTTAA-3’, and sanger-sequenced by primer 5’-TTGTAATTTTTTTTTTTATGAGTGGTTTGGTTTTA-3’. For m^5^C detection of endogenous m^5^C site in RPSA, target site was amplified using primers 5’-AAATTTTAAGAGGATTTGGGAGAAGTTTTTG-3’ and 5’-CAACCCTAAAATCAATAACCACAAAAAACCATA-3’, and sanger-sequenced by primer 5’-AAATTTTAAGAGGATTTGGGAGAAGTTTTTG-3’.

#### m^1^A detection based on mis-incorporation during reverse transcription

1 μg total RNA was reverse transcribed by HifairTM II 1st Strand cDNA Synthesis Kit (Yeasen, # 11121ES60). The region flanking m^1^A site was amplified by plasmid specific primer 5’-TTCACCGTCACCGCCGAC-3’ and 5’-CTAATTCACTCCCAACGAAGACAAGATTT-3’. The mismatch site was measured by sanger sequencing using primer 5’-CTAATTCACTCCCAACGAAGACAAGATTT-3’.

#### mRNA-BisSeq

The quality of 500 ng bisulfite-treated mRNA (see above) was assessed using Agilent RNA 6000 Pico Kit (Agilent, #NC1711873), and then subjected to NGS libraries preparation using VAHTS Stranded mRNA-seq Library Prep Kit (Vazyme). The library quality was assessed using High Sensitivity DNA Kit (Agilent, #5067-4626). Paired-end sequencing (2×150 bp) was performed with Illumina NovaSeq 6000 System by Haplox genomics center. The raw sequencing data have been deposited to GEO under the accession number GSE157368.

#### Generation of CIGAR-seq vector pool with a FURIN m^5^C reporter

A gRNA library containing 4975 gRNA targeting 829 RBP (Table EV1) was synthesized by GENEWIZ and cloned into CROP-seq vector (Addgene, #86708) at BsmBI sites. For measuring the complexity of the gRNA library, the region harboring gRNA sequence was amplified with primer pair 5’-AATGATACGGCGACCACCGAGATCTACACTCTTTCCCTACACGACGCTCTTCCGATCTCTTGTA-TATCCCTTGGAGAACCACCTTGTTG-3’ and 5’-CAAGCAGAAGACGGCATACGAGATCCACTCGTGACTGGAGTTCAGACGTGTGCTCTTCCGATCT-CGACTCGGTGCCACTTTTTCAAGTTG-3’ for NGS. Afterwards, the FURIN m^5^C reporter was amplified and used to replace WPRE cassette using ClonExpress II One Step Cloning Kit (Vazyme). During cloning of CIGAR-seq vector pool, electrocompetent Stbl3 cells (Weidi Biotechnology, CAT#: DE1046) was always used.

#### CIGAR-seq viral package

HEK293T cells were plated onto 15 cm plates at 40% confluence. The next day, cells were transfected with PEI (Polysciences, #23966-2) using 15 μg of CIGAR-seq vector, 15 μg of psPAX2 (Addgene, #12259) and 22.5 μg of pMD2.G (Addgene, #12259). Supernatant containing viral particles were harvested at 48 h and 96 h, and purified with 0.45 μm filter.

#### Genetic screen for novel m^5^C regulators

2×10^8^ HAP1 cells were infected with CIGAR-seq viral particles (MOI = 0.3) and treated with 1 μg/ml of Puromycin for 24 h post infection. Puromycin resistant cells were cultured for additional seven days, and then 2×10^8^ HAP1 cells were collected for RNA extraction. 100 ng of bisulfite-treated mRNA (see above) was quantified using Agilent RNA 6000 Pico Kit (Agilent, #NC1711873) and reverse transcribed into cDNA using HiScript II Q Select RT SuperMix (Vazyme, #R233-01). Finally, using the cDNA, CIGAR-seq NGS library was amplified with forward primer 5’-AATGATACGGCGACCACCGAGATCTACACTCTTTCCCTACACGACGCTCTTCCGATCT-GGGTTGGTTTAGGAGATATTTGAGGG-3’ and reverse primer 5’-CAAGCAGAAGACGGCATACGAGATCGTGATGTGACTGGAGTTCAGACGTGTGCTCTTCCGATCT-AACAATCCTAATACTCAAAAAAAAAAACACCA-3’. Paired-end sequencing (2×150 bp) was performed with Illumina NovaSeq 6000 System by Haplox genomics center. The raw sequencing data have been deposited to GEO under the accession number GSE157368.

#### Western blotting

Transfected HAP1 cells were collected and lysed by RIPA buffer (150 mM NaCl, 50 mM Tris, 1% EDTA, 1% NP40, 0.1% SDS). Lysate was incubated at 4 °C for 30 min, then sonicated with 10 cycles (30 s On /30 s Off), and then centrifuged at 15,000 g for 15 min at 4 °C. The total protein concentration was measured by BCA (Beyotime, #P0011). 60 μg total protein was loaded and separated on the 10% SDS-polyacrylamide gel. The protein on the gel was transfected to the polyvinylidene difluoride membranes (Immobilon-P, #IPVH00010). The membrane was incubated with primary antibody and horseradish peroxidase–conjugated secondary antibody, and then proteins were detected using the Pierce™ ECL Western Blotting Substrate (Thermo, #32209) by BIO-RAD ChemiDoc™ XRS+ system. The following antibodies were used for western blotting: NSUN2 (Proteintech, #20854-1-AP), NSUN6 (Proteintech, #17240-1-AP) and GAPDH (TransGen Biotech, #N10404).

### Computational Methods

#### CIGRA-seq data analysis

CIGRA-seq NGS data consists of paired end reads. Read1 contains the sequence of m^5^C reporter site while read2 consists of the gRNA sequence. Raw fastq data were first trimmed using fastp (Chen, Zhou et al., 2018) to remove low-quality bases (-A -w 12 --length_required 30 -q 30). Then the clean read pairs were parsed using a custom script based on pysam package. Specifically, gRNA sequence in read2 was extracted by regex module using regular expression ((CAACTTAACTCTTAAAC[ATCG]{20}CA){s<=1}). m^5^C reporter sequence was extracted in the similar way ((GTTATTT[TC]{1}TTTAAGG){s<=1}). At most 1 substitution was allowed during the pattern searching. Read pairs with both reads containing the matched pattern sequences and the m^5^C sites being C or T were kept for further analysis. Then for each gRNA sequence, the number of supported reads with reporter site being C (m^5^C) or T were calculated, and the number of C reads divided by the sum of C and T reads represented the m^5^C level. Only the extracted gRNA sequences that match exactly with the RBP gRNA sequences (Table EV1) were kept for further analysis.

To identify the high-confident candidate genes that regulate m^5^C level, information of multiple gRNAs of the same genes were combined using the Stouffer’s method. gRNAs with no more than 20 supported reads were filtered out. Genes with only one gRNA detected were filtered out. Then, given a gene i, the m^5^C level of reporter site correspondence to gRNA j is X_i,j_., m^5^C level was converted to Z-score and p value P_i,j_ was calculated under normal distribution assumption. Then a combined p value for each gene P_i_ was obtain using the weighted version of Stouffer’s method, with the logarithmic scale of read count as weight for each gRNA. Finally, P values of multiple tests were adjusted with Benjamini & Hochberg’s method.

#### mRNA-BisSeq data analysis

mRNA-BisSeq data generated in this study were analyzed following the RNA-m^5^C pipeline (Huang et al., 2019) (https://github.com/SYSU-zhanglab/RNA-m5C). Reference genomes (GRCh38) and gene annotation GTF file was downloaded from Ensemble (http://www.ensembl.org/info/data/ftp/index.html). Briefly, raw paired-end reads were trimmed using cutadapt (Martin, 2011) (-a AGATCGGAAGAGCACACGTC -A AGATCGGAAGAGCGTCGTGT -j 12 -e 0.25 -q 30 --trim-n) and then Trimmomatic(Bolger, Lohse et al., 2014) (SLIDINGWINDOW:4:25 AVGQUAL:30 MINLEN:36). Clean read pairs were aligned to both C-to-T and G-to-A converted reference genomes by HISAT2 (Kim, Paggi et al., 2019). Unmapped and multiple mapped reads were then aligned to C-to-T converted transcriptome by Bowtie2 (Langmead & Salzberg, 2012), and the transcript coordinates were liftovered to the genomic coordinates. Reads from HISAT2 and Bowite2 mapping were merged and filtered using the same criteria as in RNA-m^5^C pipeline. Bam file was transformed into pileup file (--trim-head 6 --trim-tail 6). Putative m^5^C sites were called using script m^5^C_caller_multiple.py inside RNA-m^5^C pipeline (with parameters -P 8 -c 20 -C 2 -r 0.05 -p 0.05 --method binomial). Default parameters of RNA-m^5^C scripts were used unless otherwise specified.

#### NSUN6-dependent m^5^C sites

First, to determine a set of high-confident m^5^C sites in HAP1 cells, five replicates of mRNA-BisSeq data generated from the WT HAP1 cells were used. The criteria to determine the high-confident m^5^C sites were: (1) coverage of the site being at least 20 reads in all five replicates; (2) number of reads containing the unmodified C being at least 2 in all five replicates; (3) the WT methylation level (the minimum methylation level from the five replicates) being at least 0.05. Then, to determine the NSUN6-dependent m^5^C sites, m^5^C level of the sites were at least 0.05 in WT HAP1 cells and less than 0.02 or 10% of the WT m^5^C level in NSUN6-KO HAP1 cells. NSUN2-dependent sites were defined based on the same criteria.

#### Features of the m^5^C sites

The upstream and downstream 10 bp sequences flanking the m^5^C sites were extracted from the genome. Motif analysis was performed Using ggseqlogo (Wagih, 2017) R package.

## Acknowledgments

This work was supported by the Shenzhen-Hong Kong Institute of Brain Science-Shenzhen Fundamental Research Institutions (Grant No. 2019SHIBS0002), Shenzhen Science and Technology Program (Grant No. KQTD20180411143432337, JCYJ20190809154407564 and JCYJ20180504165804015) and the National Natural Science Foundation of China (Grant No. 31701237, 31900431 and 31970601). The authors acknowledge the Center for Computational Science and Engineering of SUSTech for the support on computational resource and acknowledge the SUSTech Core Research Facilities and Guixin Ruan for technical support.

## Author contributions

W.C. and L.F. developed the concept of the project. W.W., L.F., L.Z., D.G., J.Y. and Y.T. designed and performed experiments. G.L. performed bioinformatic analysis. W.Z., M.Z., D.D., Z.S., Q.Z. and Z.D. assisted in performing experiments. W.C., L.F., G.L., Y.H., W.W., J.L., H.C. and W.S. reviewed and discussed results. W.C., L.F., G.L. and W.W. wrote the manuscript.

## Figure Legends of supplementary figures

**Figure EV1.**
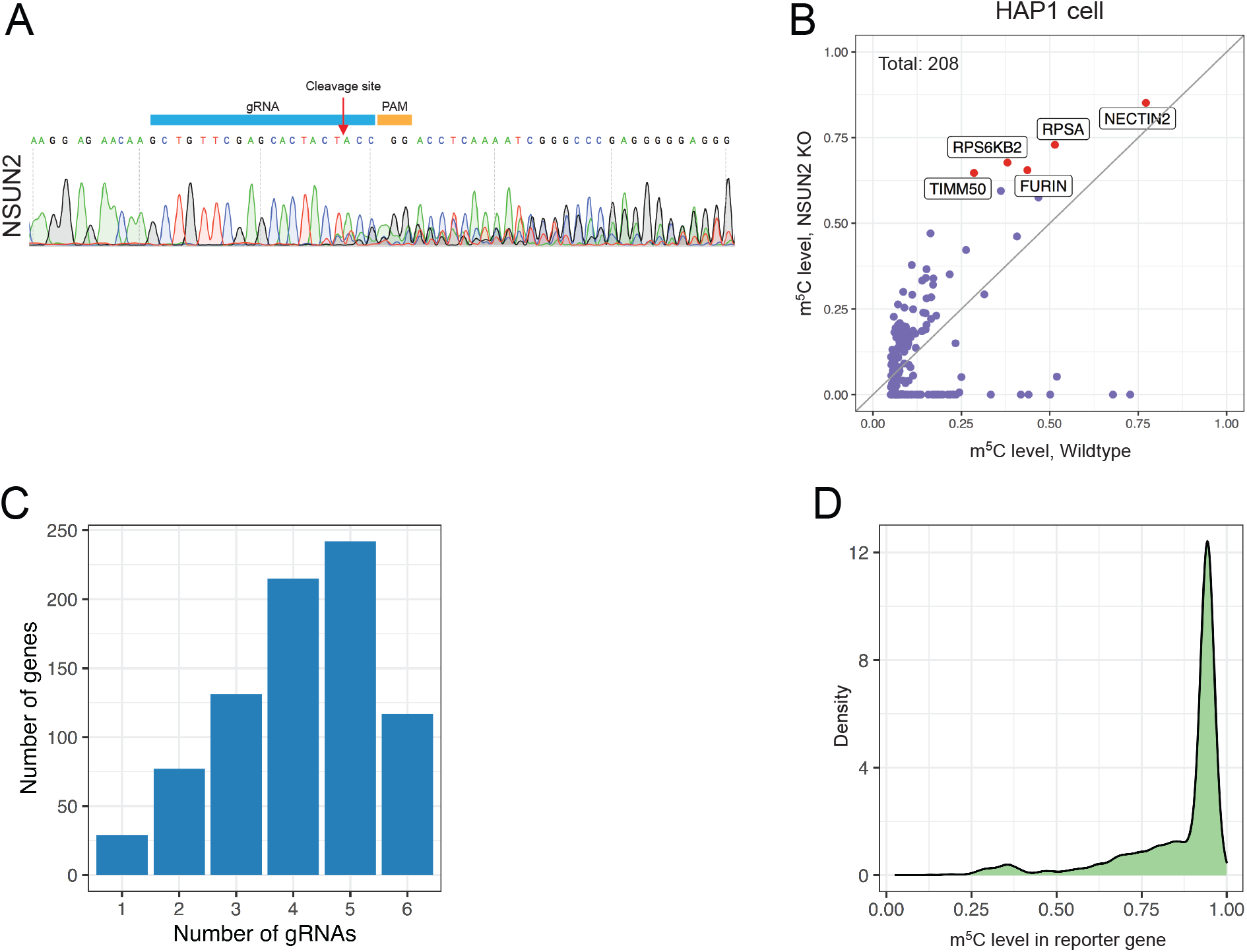
NSUN2 Knock out and the effect on the reporter as well as endogenous m^5^C sites. **A**, The NSUN2 KO efficiency in Cas9-expressing HAP1 cells. Seven days after viral transduction, NSUN2 was efficiently mutated. **B**, Scatter plot demonstrating the effect of NSUN2 knocked out in HAP1 cells. 208 m^5^C sites were identified in wildtype HAP1 cells, 90 (43.3%) of which showed significantly reduced m^5^C level in NSUN2-KO cells. **C**, Distribution of genes with different number of gRNAs detected. **D**, Distribution of m^5^C level of the reporter site associated with individual gRNA.

**Figure EV2.**
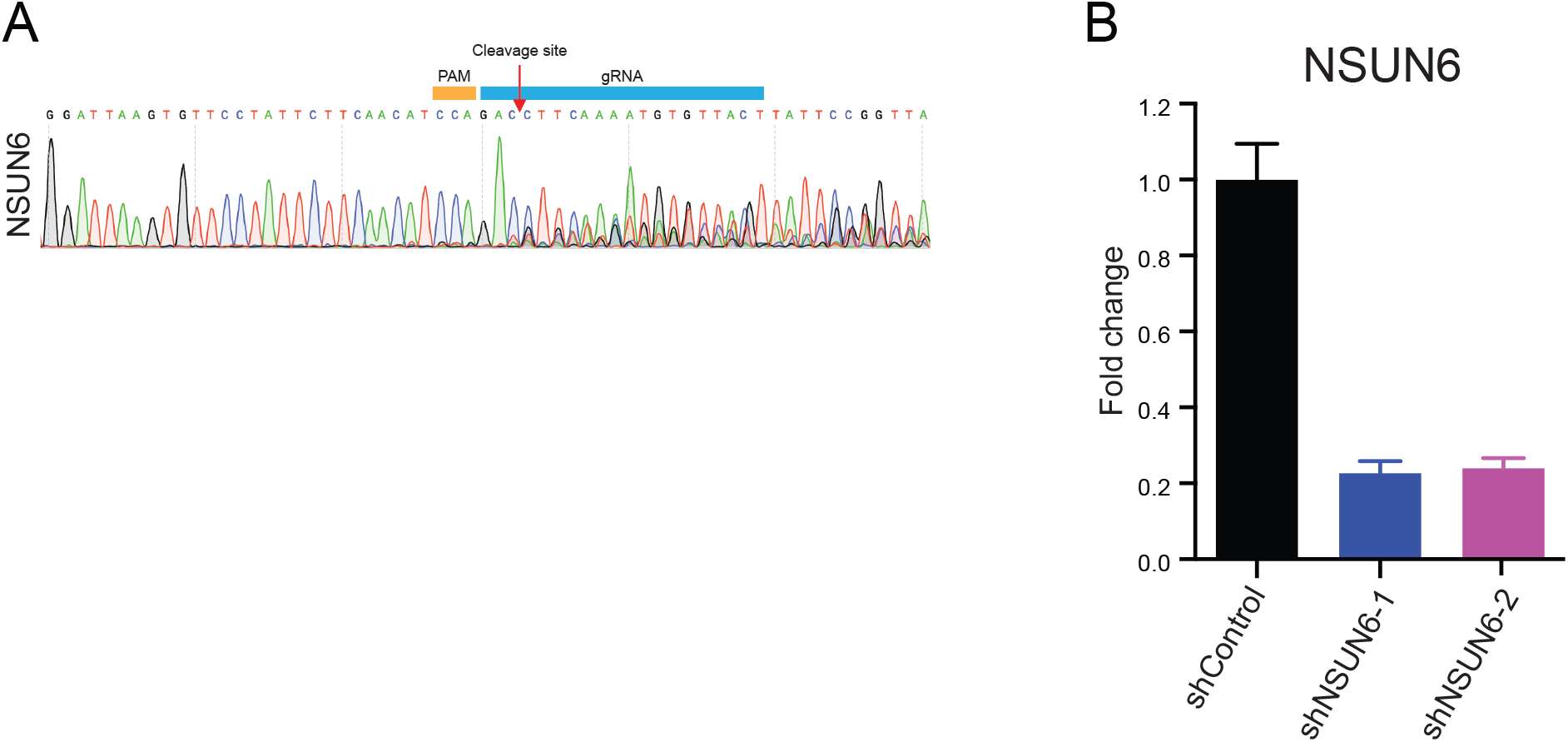
NSUN6 was efficiently knocked out or knocked down. **A**, The KO efficiency of NSUN6 in Cas9-expressing HAP1 cells. Seven days after viral transduction, NSUN6 was efficiently mutated. **B**, The knockdown efficiency of NSUN6 in HAP1 cells.

**Figure EV3.**
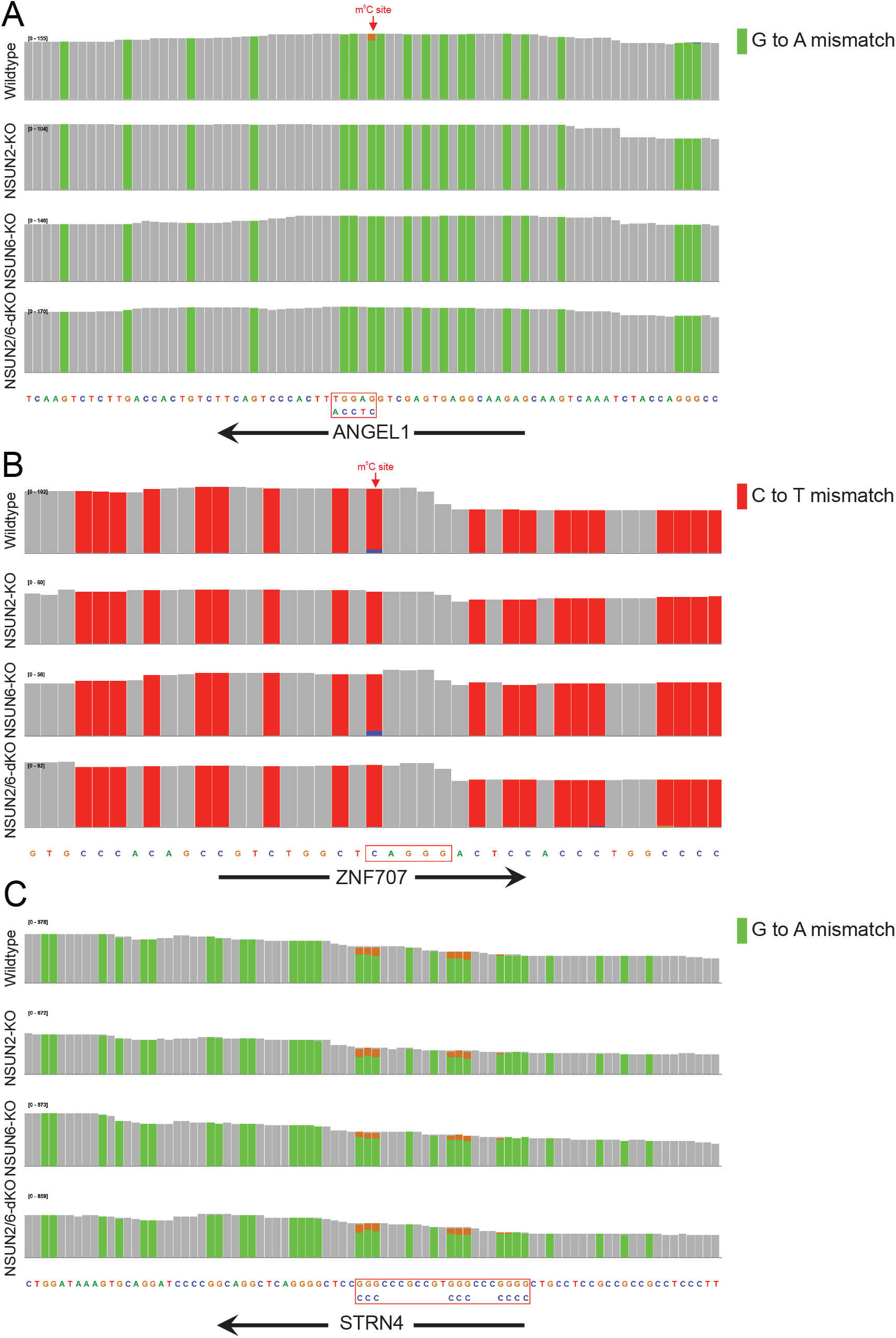
mRNA-BisSeq profiles of the three m^5^C modification sites depend on both NSUN6 and NSUN2. IGV plots showing the m^5^C sites in ANGEL1, ZNF707 and STRN4 genes. The m^5^C sites in ANGEL1 (**A**) and ZNF707 (**B**) possessed a 3’ TCCA and a 3’ AGGG motif, respectively, while the m^5^C site in STRN4 was among a cluster of “pseudo” m^5^C sites (**C**).

**Figure EV4.**
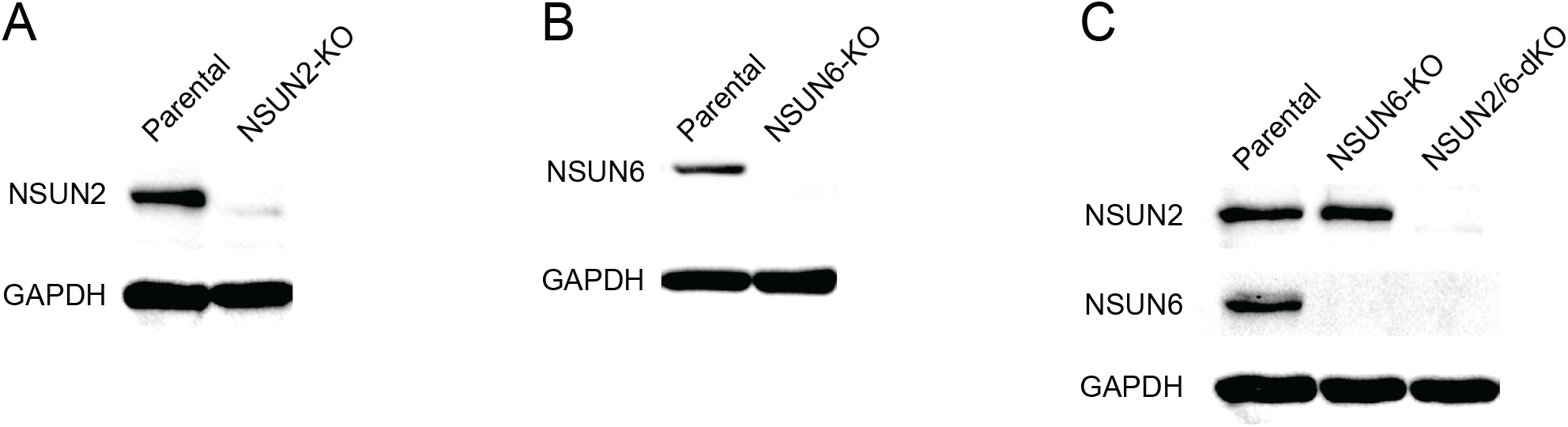
KO of NSUN2 and/or NSUN6 in HAP1 cells. Western Blot demonstrating the effect NSUN2 (**A**), NSUN6 (**B**) as well as NSUN2/6 double knockout (**C**). NSUN2/6 double knockout (NSUN2/6-dKO) HAP1 cells were established based on clonal NSUN6-KO cells.

