## Supplemental Table2 for "CIGAR-seq, a CRISPR/Cas-based method for unbiased screening of novel mRNA modification regulators"

| ID | position | gene | sequences |
|---|---|---|---|
| 1 | 11@16756661@+ | C11orf58 | cacacctataatcccagcactttgggaggccgaggtg |
| 2 | 11@66687583@- | SPTBN2 | GCATTGCAGATGGGGCCGGGCCCCGAGTCCGGGTC |
| 3 | 11@767963@- | GATD1 | gggttcaagcgattctcctgcctcagcctcccaag |
| 4 | 11@767969@- | GATD1 | gggttcaagcgattctcctgcctcagcctcccaag |
| 5 | 11@768098@- | GATD1 | cctcaggtgatccacctgcctcagcctcccaaagtg |
| 6 | 11@768104@- | GATD1 | cctcaggtgatccacctgcctcagcctcccaaagtg |
| 7 | 12@110281749@+ | ATP2A2 | CCCGCGGCCGgagtgcgaggcggaggcgaggaggccgcg |
| 8 | 12@110281750@+ | ATP2A2 | CCCGCGGCCGgagtgcgaggcggaggcgaggaggccgcg |
| 9 | 12@110281752@+ | ATP2A2 | CCCGCGGCCGgagtgcgaggcggaggcgaggaggccgcg |
| 10 | 14@74736311@+ | FCF1 | cccagccactggggaggctgaggcaggagaacctctt |
| 11 | 14@74736317@+ | FCF1 | cccagccactggggaggctgaggcaggagaacctctt |
| 12 | 15@65150178@- | CLPX | gcgattctcctgcctcagcctcccgagtagctgggat |
| 13 | 15@85234697@- | AC044860.1 | CAGCCACCCCGCGCACCGGCTACAAGCCGCCCGGGGG |
| 14 | 16@1964562@- | RPS2 | CCGCGGAAGCCACCGCGGTTCCCCATCCCAGGGCC |
| 15 | 16@2972795@+ | PAQR4 | AGGAAGGGCCATGTTGCCACATGAGCAAGCTTGGGTG |
| 16 | 16@50368785@- | BRD7 | GCCCATGTCCGAccgggccccggtgcccgcccccg |
| 17 | 18@31685103@- | | ATGCCGCGCCGGGCCCCGGCGCCCCGCGGCCGCCGCG |
| 18 | 19@13118331@+ | NACC1 | CGCCGCGGCTGCCGCTGCTgctgaggcggaggccgcggaggccg |
| 19 | 19@13118340@+ | NACC1 | CGCCGCGGCTGCCGCTGCTgctgaggcggaggccgcggaggccg |
| 20 | 1@150308596@+ | MRPS21 | cccaacactttgggaggctgaggcgggcggctcact |
| 21 | 1@156676704@- | NES | GGTAGCCGCGCACTGCCCCGCGCCACGCCTCGCCCA |
| 22 | 1@31938230@- | PTP4A2 | tcctgctgccgccgctgcctccgctgccgccgctg |
| 23 | 20@36891800@- | SAMHD1 | gttcgagcaatcctcctgcctcagcctcccaagtagc |
| 24 | 20@43667277@+ | MYBL2 | CGCGGCGCGGTCCGGGCCGGGGGGATGTCTCGGCGGA |
| 25 | 22@49961503@+ | PIM3 | GGGCGCGGCGGGCGGCGCGCGCGGCGTCATCCGCCT |
| 26 | 2@101253124@+ | CNOT11 | GGGTCCGGGAGCGGAGGCCCGGGGGGCCCCGCGGG |
| 27 | 2@216633613@+ | IGFBP2 | ctgggcgcgagtggcggcggcggcgggggcgcgcgCG |
| 28 | 2@216633616@+ | IGFBP2 | ctgggcgcgagtggcggcggcggcgggggcgcgcgCG |

| ID | position | gene | sequences |
|---|---|---|---|
| 29 | 2@216633619@+ | IGFBP2 | 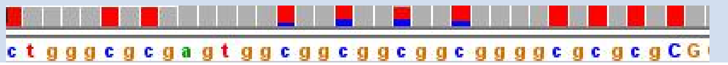  |
| 30 | 5@180071683@- | RNF130 | 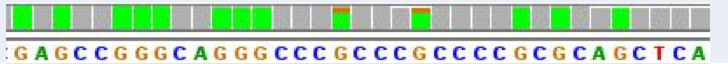  |
| 31 | 5@180071687@- | RNF130 | 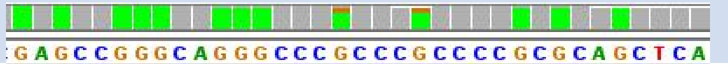  |
| 32 | 6@113859942@+ | MARCKS | 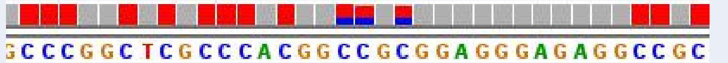  |
| 33 | 6@113859943@+ | MARCKS | 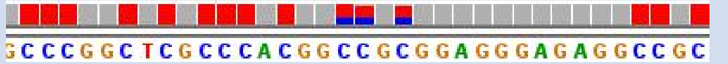  |
| 34 | 6@113859945@+ | MARCKS | 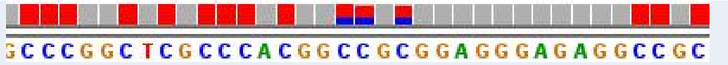  |
| 35 | 6@149724702@- | NUP43 | 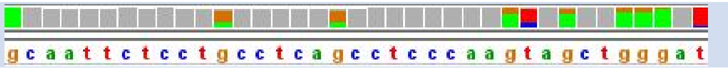  |
| 36 | 7@100070158@- | ZNF3 | 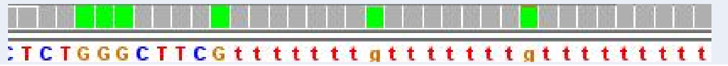  |
| 37 | 7@100070166@- | ZNF3 | 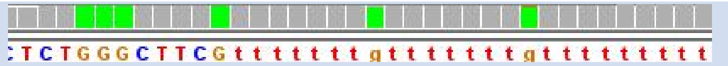  |
| 38 | X@119538511@- | CXorf56 | 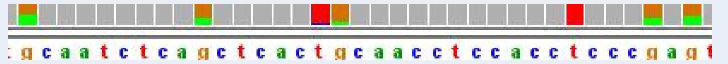  |
| 39 | X@119538527@- | CXorf56 | 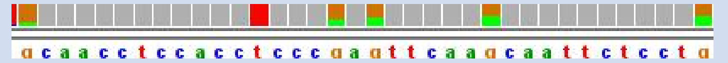  |
| 40 | 7@151080695@- | FASTK | 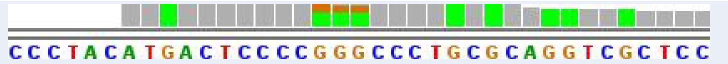  |
| 41 | 7@151080696@- | FASTK | 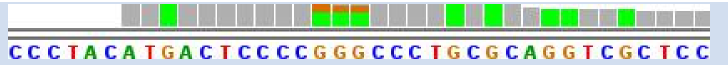  |
| 42 | 7@151080697@- | FASTK | 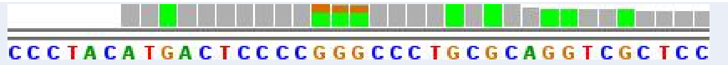  |
| 43 | 8@12755322@- | LONRF1 | 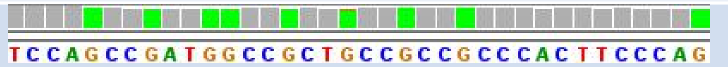  |
| 44 | 8@17156688@+ | ZDHHC2 | 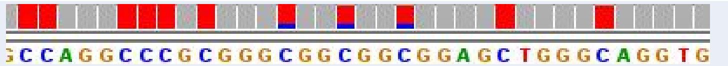  |
| 45 | 8@17156691@+ | ZDHHC2 | 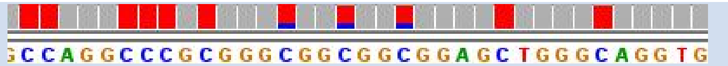  |
| 46 | 8@17156694@+ | ZDHHC2 | 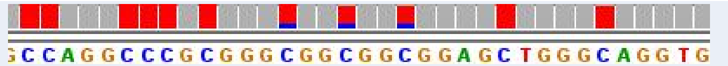  |
| 47 | MT@14423@- | MT-ND6 | 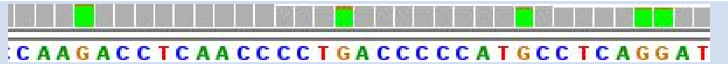  |
| 48 | MT@1486@+ | MT-RNR1 | 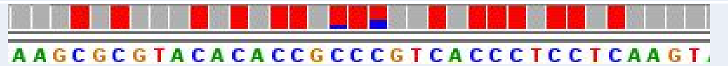  |
| 49 | MT@1488@+ | MT-RNR1 | 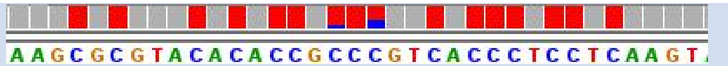  |
| 50 | MT@8387@- | | 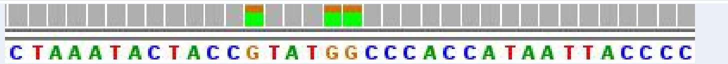  |
| 51 | MT@8391@- | | 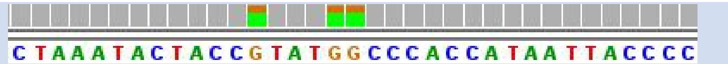  |
| 52 | MT@8392@- | | 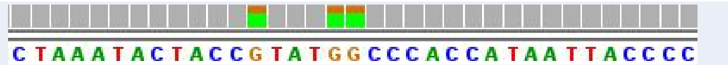  |
| 53 | 6@42989628@+ | **PPP2R5D** | 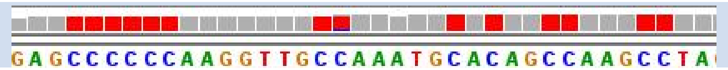  |
| 54 | 7@134293043@- | **SLC35B4** | 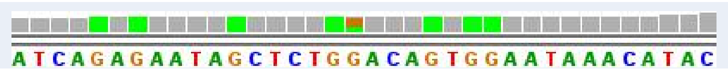  |
| 55 | 5@179616893@- | **HNRNPH1** | 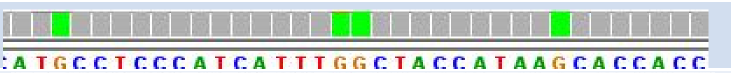  |
| 56 | 5@80626854@- | **DHFR** | 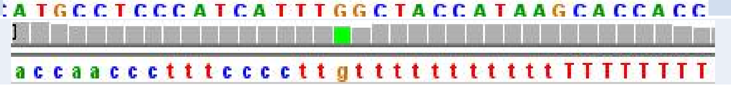  |
